# Transposon Mutagenesis Reveals RBMS3 as a Promoter of Malignant Progression of BRAF^V600E^-Driven Lung Tumorigenesis

**DOI:** 10.1101/2022.02.28.482366

**Authors:** Aria Vaishnavi, Joseph Juan, Michael T. Scherzer, J. Edward VanVeen, Christopher Stehn, Christopher S. Hackett, Adam Dupuy, Steven A. Chmura, Louise van der Weyden, Justin Y. Newberg, Karen M. Mann, Annie Liu, Alistair G. Rust, William A. Weiss, David J. Adams, Allie Grossmann, Michael B. Mann, Martin McMahon

## Abstract

Mutationally-activated BRAF^V600E^ is detected in ~2% of all human non-small cell lung cancers (NSCLC), and serves as a predictive biomarker for treatment of patients with FDA-approved pathway-targeted therapies that inhibit signaling by the BRAF^V600E^ oncoprotein kinase. In genetically engineered mouse (GEM) models, expression of BRAF^V600E^ in alveolar type 2 (AT2) pneumocytes initiates the development of benign lung tumors that, without additional genetic alterations, rarely progress to malignant lung adenocarcinomas. To identify genes that might cooperate with BRAF^V600E^ for malignant lung cancer progression we employed *Sleeping Beauty* (*SB*)-mediated transposon mutagenesis, which dramatically accelerated the onset of lethal lung adenocarcinomas. Amongst the diverse group of genes identified by this *in vivo* screen was *Rbms3* (<u>R</u>NA <u>b</u>inding <u>m</u>otif <u>s</u>ingle-stranded interacting protein <u>3</u>), an RNA-binding protein implicated as a possible tumor suppressor. Using CRISPR/CAS9 gene editing we confirmed that RBMS3 silencing cooperated with BRAF^V600E^ to promote progression of malignant lung cancer with a distinct micropapillary architecture. Moreover, RBMS3 silencing also cooperated with BRAF^V600E^ to promote the growth of lung organoids *in vitro*. BRAF^V600E^/RBMS3^Null^ lung tumors displayed elevated expression of b-catenin (CTNNB1), suggesting that RBMS3 silencing may result in elevated signaling through the WNT>CTNNB1>c-MYC pathway. Finally, analyses of patient samples in The Cancer Genome Atlas (TCGA) revealed that the region of chromosome 3 encompassing *RBMS3* is frequently lost in NSCLC and correlates with poor patient prognosis. Collectively, *SB*-mediated transposon mutagenesis has revealed the ability of a novel tumor suppressor, *RBMS3*, to cooperate with BRAF^V600E^ to promote lung carcinogenesis, and suggests that RBMS3 silencing may contribute to malignant progression of numerous human lung cancers.

**SIGNIFICANCE:** The BRAF^V600E^ oncoprotein kinase is a potent initiator of benign lung tumorigenesis, but is insufficient to elicit malignant lung adenocarcinoma without additional cooperating alterations. *Sleeping Beauty*-mediated transposon mutagenesis has revealed a number of genes that cooperate with BRAF^V600E^ to promote lung cancer progression, in particular *Rbms3*, which encodes an RNA binding protein. Hence, this genetic screen provides a deeper understanding of the molecular mechanisms underlying BRAF^V600E^-driven lung carcinogenesis, and is an important step improving our ability to successfully target this disease.

## INTRODUCTION

Non-small cell lung cancer (NSCLC) is the leading cause of cancer-related death globally, with adenocarcinoma being the singe largest sub-type (Siegel et al. 2016). Over the past decade, treatment outcomes have improved for lung cancer patients whose tumors are driven by actionable oncogenic mutations. For example, mutational activation of genes such as *Epidermal Growth Factor Receptor* (*EGFR*), *Anaplastic Lymphoma Kinase* (*ALK)* or *Neurotrophin Tyrosine Kinase Receptor Type 1* (*NTRK1)* serve as predictive biomarkers for the use of FDA-approved targeted drugs osimertinib, alectinib or larotrectinib, respectively (Peters et al. 2017; Drilon et al. 2018; Soria et al. 2018). However, our understanding of how such oncoproteins promote the initiation, progression and maintanence of lung adenocarcinoma remains incomplete. Moreover, since most lung cancers result from the accrual of multiple genetic/epigenetic alterations that cooperate in the conversion of normal lung cells into malignant lung cancer cells, we need a deeper mechanistic understanding of how such cooperation operates at the molecular and cellular levels and how it may influence lung cancer therapeutic strategies.

One of the actionable mutations in lung cancer is *BRAF*^*T1799A*^, that is detected in 2-4% of patients, which translates to ~3300 patients per year in the US (Heist and Engelman 2012; Vultur and Herlyn 2013). BRAF plays an important role in the activation of the RAS-regulated RAF>MEK>ERK MAPK signal transduction pathway, a pathway of critical importance in normal development and tissue homeostasis, which is frequently dysregulated in human tumorigenesis. *BRAF*^*T1799A*^ encodes BRAF^V600E^, a constitutively active oncoprotein kinase, which is mutated in numerous malignancies including melanoma, hairy cell leukemia, colorectal, pancreatic and thyroid cancers (Cancer Genome Atlas Research 2014). The importance of BRAF^V600E^ in cancer maintenance is emphasized by FDA approval of three pairwise targeted therapeutic combinations (1. Vemurafenib & cobimetinib; 2. Dabrafenib & trametinib and; 3. Encorafenib & binimetinib) that target BRAF^V600E^>MEK>ERK signaling and most importantly are superior to the standard of care therapy (Larkin et al. 2014; Planchard et al. 2017; Dummer et al. 2018). However, although responses to vertical inhibition of BRAF^V600E^ signaling often elicit striking responses, most patients eventually develop lethal drug resistant disease, leaving additional room for improvement to these targeted therapeutic approaches.

Genetically engineered mouse (GEM) models of human cancer provide a powerful platform to test hypotheses regarding tumor initiation, cancer progression and maintenance. *Braf*^*CA*^, as well as *Braf*^*CAT*^mice were engineered to express normal BRAF prior to CRE-mediated recombination after which BRAF^V637E^ (analogous to human BRAF^V600E^, the nomenclature that we will use to describe this model hereafter) is expressed in cells in a temporally and spatially restricted manner (Dankort et al. 2007; van Veen et al. 2016; van Veen et al. 2019). The *Braf*^*CAT*^ mice were further adapted to express the tdTomato flourescent reporter upon additional of CRE as well. Expression of BRAF^V600E^ in the distal epithelium of the mouse lung elicits clonal tumorigenic outgrowths of alveolar type 2 (AT2) pneumocytes (Dankort et al. 2007). Growth of the resulting benign lung adenomas is constrained by a senescence-like growth arrest that is triggered by an insufficiency in WNT>β-catenin>c-MYC signaling (Juan et al. 2014). However, mutationally activated BRAF^V600E^ cooperates with numerous genetic alterations including silencing of TP53 or INK4A/ARF, or expression of mutationally activated PI3-kinase-α (PIK3CA^H1047R^) for malignant lung carcinogenesis (Dankort et al. 2007; Trejo et al. 2013). *Sleeping Beauty* (SB) insertional mutagenesis has facilitated the identification of genes that participate in various aspects of tumorigenesis in GEM models, and was previously used to identify genes that cooperate with BRAF^V600E^ in melanomagenesis (Mann et al. 2015; de la Rosa et al. 2017; Montero-Conde et al. 2017; Weber et al. 2020). Here, we employed *SB*-mediated insertional mutagenesis to identifiy numerous candidate genes that cooperate with BRAF^V600E^ in lung carcinogenesis. Furthermore, CRISPR/Cas9 gene editing allowed the validation of *Rbms3* as a novel suppressor of BRAF^V600E^-driven lung cancer. Importantly, these mouse data are consistent with the observation that loss of some or all of chromosome 3p, where human *RBMS3* is located, is a common event in numerous human lung cancers. In this study, we report that this GEM approach may have helped identify to a previously underappreciated, but frequent lung cancer tumor suppressor gene that can cooperate with BRAF^V600E^ to accelerate lung cancer progression.

## RESULTS

### The <u>B</u>raf^CA^|<u>S</u>B|<u>L</u>ung (BSL) Insertional Mutagenesis Screen

To conduct the insertional mutagenesis screen, we utilized the following genetic elements: 1. *Braf*^*CA*^, the CRE-activated allele of *Braf* (Dankort et al. 2007); 2. *RCL::SB*, <u>*R*</u>*osa26*-<u>*C*</u>*AGGS*-<u>*L*</u>*SL*-<u>*SB*</u>11, a CRE-activated SB11 transgene that was recombined into the *Rosa26* locus and; 3. *C4*^*T2/Onc2*^, a *T2/Onc2* (6070) transposon donor located on chromosome 4. To peform the screen we generated two cohorts of mice (n=50/cohort): 1. <u>*B*</u>*raf*^*CA*^; <u>*R*</u>*CL*::*SB*; <u>*T*</u>2/*Onc*2 (*BRT* mice) and; 2. <u>*B*</u>*raf*^*CA*^; <u>*R*</u>*CL*::*SB* (*BR* mice), which lack the *T2/Onc2* transposon. Lung tumorigenesis was initiated by intranasal instillation of 10^6^ pfu of an adenovirus encoding CRE recombinase (Ad-CMV-CRE). In control *BR* mice, the action of CRE recombinase delivered to the lung should result in co-expression of BRAF^V600E^ plus SB11 but with no *T2/Onc2* transposon, which is only present in the *BRT* mice. Initiated mice were monitored for signs of lethal lung tumorigenesis for 250 days (Fig. 1A). As anticipated, none of the control *BR* mice had to be euthanized over the monitoring period, whereas ~70% of *BRT* mice developed end-stage disease as evidenced by labored breathing and/or loss of body weight that required euthanasia. Median survival of initiated BRT mice was 205 days compared to 338 days for the BR mice (Fig. 1A, p=0.00000383). These data suggested that *SB*-mediated mobilization of the *T2/Onc2* transposon dramatically accelerated malignant progression of BRAF^V600E^-driven lung tumorigenesis in *BRT* mice.

**Figure 1.**
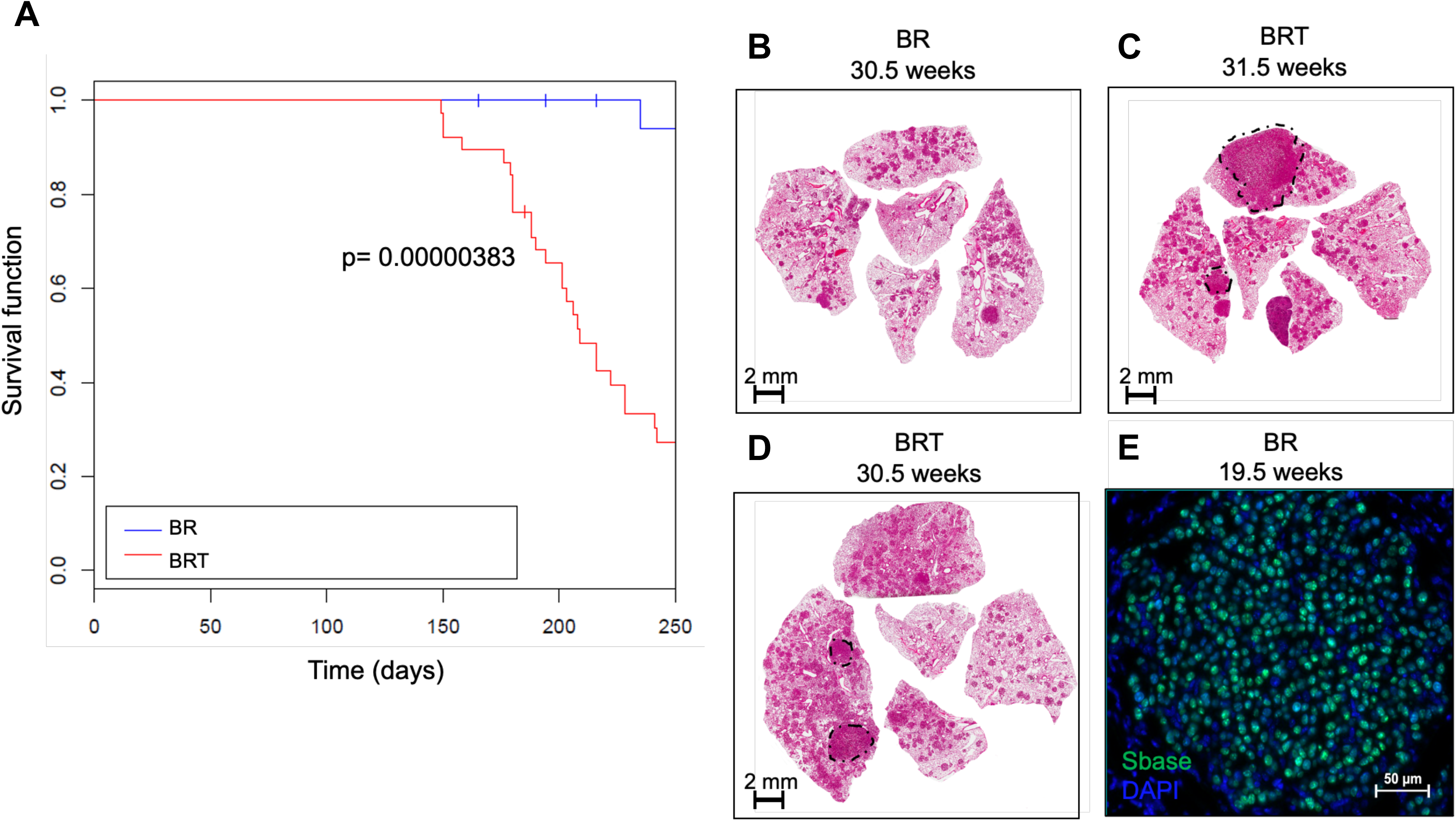
The *Sleeping Beauty* (SB) transposon system promotes lethal progression in a mouse model of BRAF^V600E^-driven lung cancer. **(A)** Kaplan meier survival analysis of 50 <u>B</u>RAF^CA^ and SB (CAGG/<u>R</u>26^LSL-SB11^) **(BR)** mice without a <u>T</u>2/Onc2 transposon or (**BRT)** mice with a donor (T2/Onc2) for 250 days. Mice were initiated through intranasal instillation with 10^6^ pfu of Ad-CMV-CRE. Statistical analysis was performed using a log-rank Mantel-Cox test where p = 0.00000383. **(B-D)** Histological analyses of formalin fixed paraffin embedded (FFPE) tumor bearing lung sections stained with hematoxylin and eosin (H&E). **(E)** Expression of SB transposase in **BRT** or **BR** mouse lung tumors at 19.5 weeks through immunofluorescence analysis of FFPE lung histological sections. DNA (DAPI) in blue, SB Transposase in green.

At euthanasia, all mice were subjected to necropsy revealing that the lungs of BRT mice displayed histological evidence of malignant lung cancer *in situ* whereas the lungs of BR mice largely contained benign adenomas (Fig. 1B). BRT mice developed a wide range of tumor grades from benign adenomas to adenocarcinomas (Fig. 1B, 1C, 1D). To identify sites of *T2/Onc2* insertion in the mouse genome, formalin-fixed, paraffin-embedded (FFPE) tissue from 28 individual lung cancers were microdissected from H&E-stained sections. Tumors from within the same mouse were selected for microdissection based on higher tumor grade (adenocarcinoma), larger tumor size, and detection of the *SB* transposase by immunofluorescences (Fig. 1E). Genomic DNA isolated from the tumors was used in splinkerette PCR (sPCR) to amplify the chromosomal *T2/Onc2* insertion sites by splinkerette PCR (sPCR) (**Fig. 1C**). Tumors were prioritized and selected for sequencing analysis, based on higher grade (adenocarcinomas) and larger size (indicative of progression past growth arrest seen in adenomas)(Su et al. 2008). DNA sequences corresponding to mouse genomic regions flanking *T2/Onc2* insertions sites were mapped using complimentary standard bioinformatics approaches including the locus-centric Gaussian kernel convolution, as well as gene-centric common insertion site (gCIS) analysis, following by SB Driver Analysis (Brett et al. 2011; Newberg et al. 2018a). These methods uniformly help identify genomic regions with a higher density of transposon insertions, and strongly suggest these regions contain a potential cancer-relevant gene.

Collectively, this pool of bioinformatics analyses, tailored strategically towards a *SB* mutagenesis screen, resulted in a stratified list of genes that cooperated with BRAF^V600E^ to promote lung tumor progression in a statistically robust maner (Fig. 2 and Table 1). This collection of statistically significant candidate driver genes include predominantly inactivating events, likely observed in potential tumor suppressor genes based on bioinformatics analysis of transposon insertion sites (Table 1; p < 0.005). It is important to clarify that in the study reported here, the term “driver” with regard to candidate genes identified in this screen, does not refer to oncogenic driver, as is more commonly observed in the literature, but rather that it was implicated as important in the SB screen. Included in this list were several previously implicated tumor suppressor genes, such as *Cux1* and *Rbms3*, but also identified, as well as *Gnaq*, a recurrent oncogenic alteration seen in uveal melanoma (Ramdzan and Nepveu 2014; Liang et al. 2015; Truong et al. 2020). In addition, the tumorigenic role of some of the genes we identified are likely to be context-dependent, based on both the cell type and their function, such as *Snd1* and *Foxp1* (Koon et al. 2007; Chidambaranathan-Reghupaty et al. 2018). Ultimately, our SB screen was able to successfully identify a wide variety of potential promoters of BRAF^V600E^-driven lung cancer progression.

**Table 1.** SB Trunk Driver genes involved in lung adenocarcinoma progression of BRAF^V600E^-initiated tumors. CIS genes containing 3 or more sequence reads per insertion from 3 or more tumors and have multiple testing corrected p < 0.05 by SB Driver Analysis.

| Gene | p value (adj) |
| --- | --- |
| *Cux1* | 3.25E-99 |
| *Wapal* | 1.24E-79 |
| *Top1* | 1.50E-71 |
| *Cmip* | 5.90E-50 |
| *Gnaq* | 5.74E-23 |
| *Snd1* | 3.75E-14 |
| *Foxp1* | 1.73E-11 |
| *Rbms3* | 4.99E-03 |

**Figure 2.**
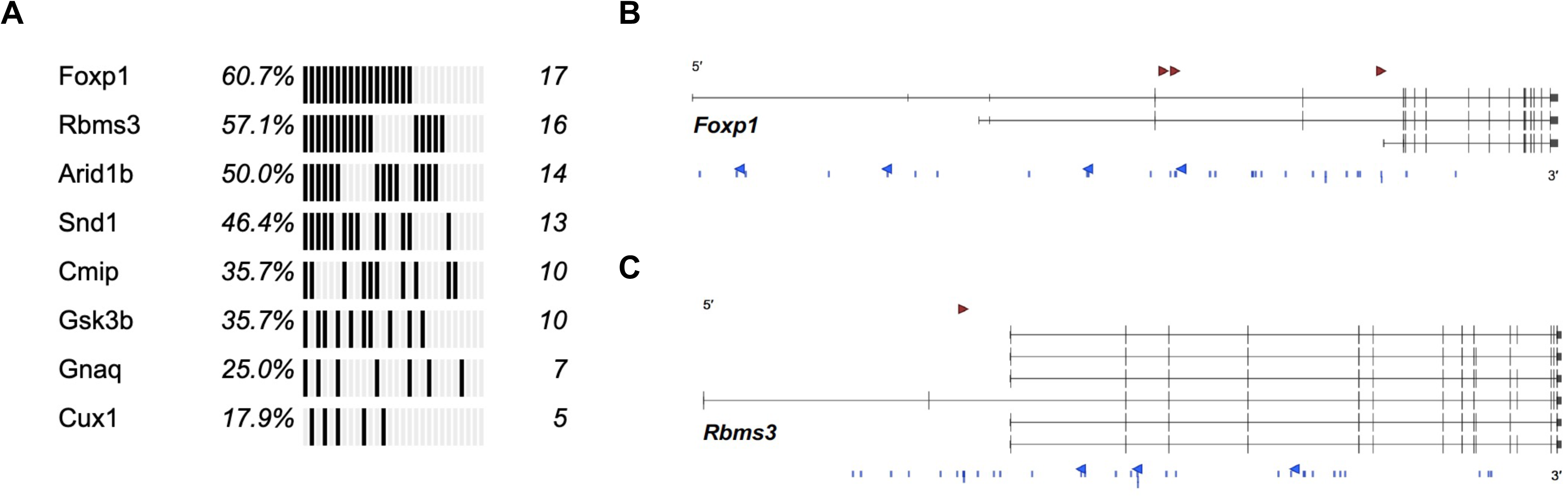
Genomic landscape of *SB*|*Braf* lung drivers. **(A)** Oncoprint of statistically significant drivers in the 28 SB driver genes occurring in oncogenic *Braf* lung tumors detected using SB Trunk Drvier analysis, from SB common integration regions (CIRs) and unique, directional SB insertions at TA-dinucleotides. **(B)** SB insertions at TA-dinucleotides with sense (red arrowhead) and anti-sense (blue arrowheads) SB insertions, and within CIRs (blue lines), for gene *Foxp1* (3 transcripts of the genes are shown). **(c)** SB insertions at TA-dinucleotides with sense (red arrowhead) and anti-sense (blue arrowheads) SB insertions, and within CIRs (blue lines), for genes *Rbms3* (6 transcripts of the candidate gene are shown).

### Rbms3 is a tumor suppressor that cooperates with BRAF^V600E^ in lung carcinogenesis

The two most significantly enriched candidate genes in this screen were *Foxp1* and *Rbms3*, which carried 17 and 16 SB insertions, respectively, in a manner consistent with gene inactivation suggesting that both might be tumor suppressors (Fig. 2A–C). However, as *Foxp1* is transcriptional regulator of a wide range of biological processes (Koon et al. 2007) and it can function both an oncogene and a tumor suppressor gene, depending on the context. Thus the context-dependent roles of *Foxp1* made potential validation of this candidate less straightforward than a clear tumor suppressor gene. RBMS3 is a single-stranded RNA binding protein that has been implicated as a potential tumor suppressor in several malignancies, including squamous cell lung cancer (Penkov et al. 2000; Li et al. 2011; Chen et al. 2012; Liang et al. 2015; Wu et al. 2017; Yang et al. 2017; Wu et al. 2019; Zhu et al. 2019). Furthermore, *Rbms3* has been implicated to play a role regulating WNT signaling, which is known to play critical roles in development and homeostasis of the lung, as well as the tumourigenesis of KRAS^G12D^ and BRAF^V600E^ oncogene-driven lung cancer (Desai et al. 2014; Juan et al. 2014; Kadzik et al. 2014; Tammela et al. 2017; Nabhan et al. 2018). Additionally, because the molecular and cellular mechanisms underlying Rbms3’s potential tumor suppressor role, as well as the role of RBMS3 in lung adenocarcinoma remain poorly understood, *Rbms3* stood out as a previously unknown, but important node in this disease subtype. This led us to focus our studies on validate this particular hit in BRAF^V600E^-driven lung cancer.

To validate the potential tumor suppressor activity of *Rbms3* in BRAF^V600E^-driven lung carcinogenesis, either non-targeting control or sgRNAs targeting *Rbms3* were cloned into a lenti-sgRNA-CRE vector used for *in vivo* expression of CRE recombinase and various sgRNAs (Chiou et al. 2015). To conduct this experiment, we used mice carrying our *Braf*^*CAT*^ allele or an *H11*^*LSL-CAS9*^ allele, which allows for CRE-activated Cas9 transgene expression from the *Hipp11* (*H11)* “safe-harbor” locus, either alone or in combination with one another (Chiou et al. 2015). Lentiviral supernatants introduced into the lungs of recipient mice by intratracheal intubation and the mice were euthanized at 11 weeks post-initiation for assessment of lung tumorigenesis. Infection of *Braf*^*CAT*^ mice alone with lenti-sgRbms3-CRE virus or of *Braf*^*CAT*^; *H11*^*LSL-CAS9*^ mice with lenti-sgNT-CRE virus led to the development of benign BRAF^V600E^-driven lung tumors, as expected (Figs. 3A & B). Strikingly, however, infection of *Braf*^*CAT*^; *H11*^*LSL-CAS9*^ mice with lenti-sgRbms3-CRE virus resulted in a signficiant increase in overall lung tumor burden, as well as a significant increase in tumor diameter compared to relevant controls (Fig. 3C, 3E, and 3F). By important contrast, infection of the lung epithelium of *H11*^*LSL-CAS9*^ mice alone with lenti-sgRbms3-CRE virus had no substantial effect on the lungs with no evidence of lung tumorigenesis (Fig. 3D). The observed increase in overall tumor size is important to note since it indicates that *Rbms3* silencing was sufficient to bypass the senescence-like growth arrest observed in benign mouse lung tumors driven solely by BRAF^V600E^ (Dankort et al. 2007; Juan et al. 2014). DNA isolated directly from large tumors demonstrated genomic editing in the *Rbms3* gene compared to controls (as assessed using the Surveyor assay; Supplementary Fig. 1A) and tumor RNA showed a significant reduction in *Rbms3* mRNA expression compared to relevant controls (as assessed by quantitiatve RT-PCR; Supplementary Fig. 1B). Collectively, these data suggests three central conclusions: 1. *Rbms3* was appropriately edited in the *in vivo* models using CRISPR/Cas9 editing technology; 2. CRISPR/Cas9-mediated *Rbms3* gene editing resulted in substantially reduced *Rbms3* mRNA expression and; 3. silencing of *Rbms3* was sufficient to promote enhanced lung cancer disease when combined with BRAF^V600E^.

**Figure 3.**
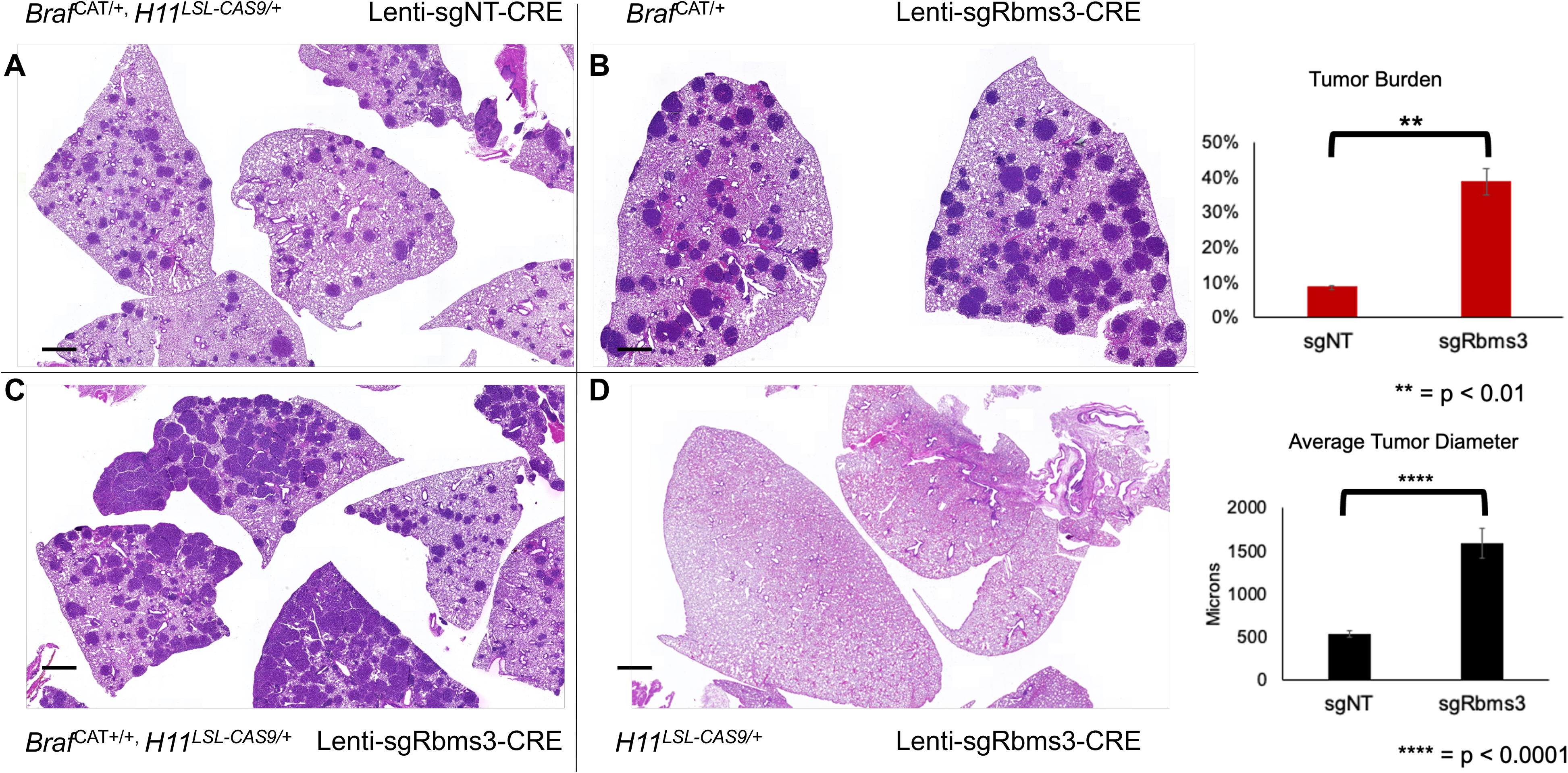
CRISPR/Cas9 disruption of RBMS3 cooperates with BRAF^V600E^ in a mouse model of lung lung cancer. (**A**) – (**D**) Panels of 4 genotypes of harvested mouse lung sections following necropsy analyses stained with hematoxylin and eosin (H&E) 13 weeks post initiation with 5 x 10^4^ pfu lenti-CRE. CRISPR/CAS9-mediated genome editing was used in panels **A**, **C**, **D** to edit *Rbms3 in vivo*. Genotype and tumor burden calculation of each experimental group was (**A**) sgNT-CRE virus in BRAF^CAT/+;^ H11^LSL-CAS9^: **8.5%**. (**B**) sgRbms3-CRE virus in BRAF^CAT/+^: **7.7%**. (**C**) sgRbms3-CRE virus in BRAF^CAT/+;^ H11^LSL-CAS9/+^: **38.8%**. (**D**) sgRbms3-CRE virus in H11^LSL-CAS9/+^: **<1%**. Black bar in bottom left of each panel represents a 1000 micron scale bar. (**E**). Quantification of tumor burden from genotypes in panel **A** compared to panel **C**. Tumor bearing lungs from panel **B** were identical to panel **A**. A paired T-test was used to determine statistical significance; p < 0.01. (**F**) Quantification of tumor diameter was performed in microns using 25 tumors from genotypes in panel **A** compared to panel **C** using the 3D Histech MIDI Slide Scanner QuantCenter. Comprehensive analyses was conducted with over 200 lung lobes. N = 50 mice individual or (biological replicates). N = 2 experimental or (technical replicates) were performed comparing the indicated genotypes in **(A)** and **(C)**. A paired T-test was used to determine statistical significance; p < 0.0001.

Previous research has established that BRAF^V600E^-driven mouse lung are benign adenomas displaying charcteristic cytomorphological features at the histological level (Dankort et al. 2007; Juan et al. 2014). Histological examination of the lung tumors in this study was consistent with these previous studies, revealing discreet papillary adenomas with a central fibrovascular core and with well-circumscribed borders (Supplementary Fig. 2A–B). Interestingly, the larger BRAF^V600E^-driven lung tumors (>1000 microns) that emerged in the context of reduced/silenced *Rbms3* expression displayed evidence of cancer progression including poorly circumscribed borders, inceased ratio of nuclear to cytoplasmic content in the tumor cells, and avascular neoplastic nests free floating in air spaces (Supplementary Fig. 2D–F). Most importantly, we observed a distinct micropapillary architecture that has previously been shown to be a strong indicator of malignant adenocarcinoma of the lung in human patients that are driven by EGFR, KRAS or BRAF oncoproteins (De Oliveira Duarte Achcar et al. 2009) (Supplementary Fig. 2E–F). This distinctive phenotype also suggests that silencing of RBMS3 in combination with BRAF^V600E^ in the mouse lungs tends to mimic key disease features of human BRAF^V600E^-driven lung adenocarcinomas. Overall, our histological evaluation of BRAF^V600E^-driven lung tumors (>1000 microns) that emerged in the context of reduced/silenced RBMS3 expression was that they displayed features of cancer progression compared to BRAF^V600E^-driven benign lung adenomas. These data are therefore consistent with the hypothesis that *Rbms3* is a tumor suppressor gene, such that it’s reduced/silenced expression promotes lung cancer progression.

As an additional approach to test oncogenic cooperation between BRAF^V600E^ expression and silencing of RBMS3 we developed a mouse lung organotypic model system. 3D organotypic culture techniques have proven highly advantageous for studying both normal and malignant cell proliferation and are an excellent complement to both in vivo studies and conventional 2D tissue culture experiments (Rock et al. 2009; Boj et al. 2015; Fatehullah et al. 2016). When digested suspensions of normal mouse lung single cells are seeded in matrigel culture in the presence of WNT signaling pathway ligands (R-Spondin1 and Noggin) and MAPK pathway growth factors (including FGF and EGF), organoids will develop over the course of 7 days. However, the requirement for EGF and FGF for lung organoid growth can be supplanted if the expression of BRAF^V600E^ is initiated in the lungs of *Braf*^*CAT*^ mice using CRE recombinase. Constitutive pathway activation of the BRAF signaling node overcomes the need for ligand-mediated pathway activation to promote lung cell proliferation and enables the selection of BRAF^V600E^-positive cells. Consequently, we isolated single-cell suspensions digested from lung tissue from *Braf*^*CAT*^; *H11*^*LSL-CAS9*^ mice infected with either control virus, lenti-sgNT-CRE, left or lenti-sgRbms3-CRE viruses. After one week of culture, organoids established from mouse lungs in which BRAF^V600E^ was expressed and *Rbms3* was silenced were substantially larger than those observed with BRAF^V600E^ expression alone (Fig. 4A). Furthermore, as expected, the organoids derived from *Braf*^*CAT*^; *H11*^*LSL-CAS9*^ mice infected with lenti-sgRbms3-CRE demonstrated lower *Rbms3* mRNA expression compared to control and also compared to cells engineered with ectopic overexpression of *Rbms3* (Fig. 4B). These data provide additional compelling evidence that *Rbms3* silencing cooperates with BRAF^V600E^ to promote lung organoid growth *in vitro* consistent with our *in vivo* validation experiments described above.

**Figure 4.**
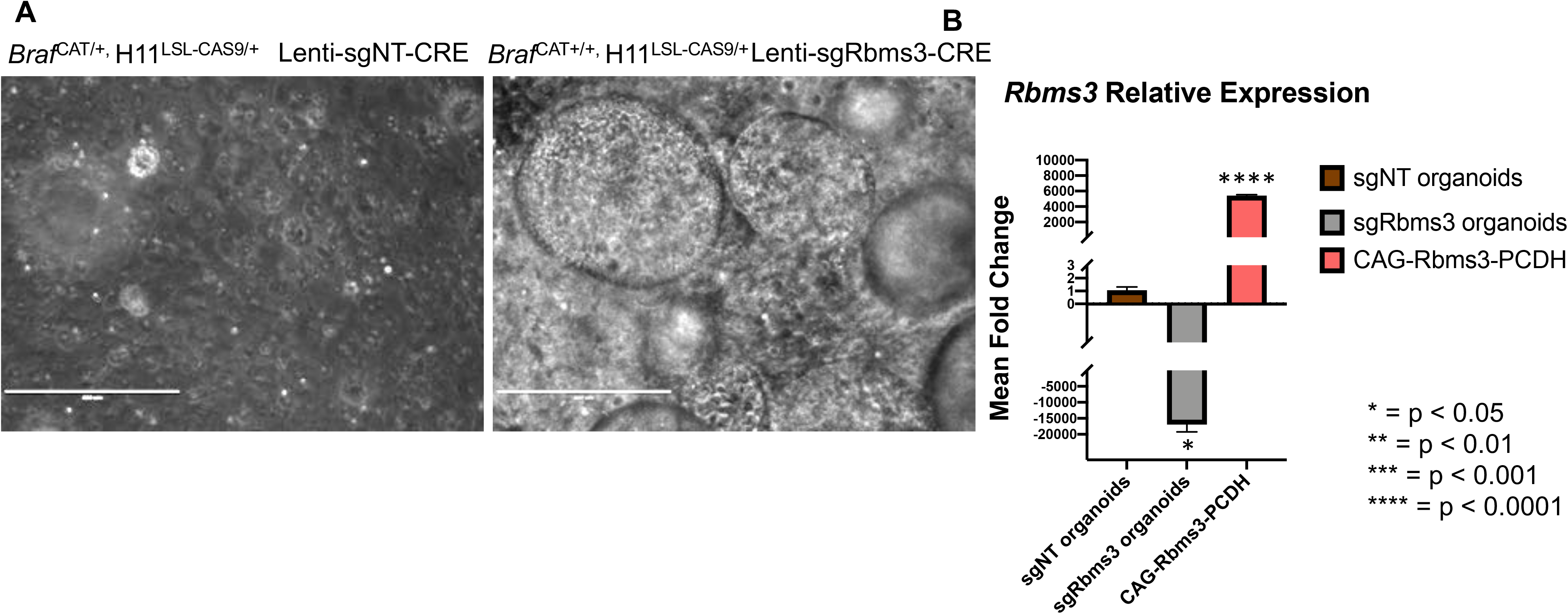
Mouse-derived organoid culture models support *Rbms3* is a cooperating gene in BRAF^V600E^-driven lung cancer. (**A**) Representative images are shown of qualitative analyses of phase contrast images of organoids established following tumor dissociation from the indicated mouse genotypes at 7 days post-initiation of organoids. White scale bar indicates 400 microns, and was taken with 10x magnification. N = 2 experimental replicates with 3 biological replicates leveraging multiple unique mouse lung lobes. (**B**). qRT-PCT analysis of *Rbms3* gene expression in organoids derived from BRAF^CAT/+;^ H11^LSL-CAS9^ mice labeled by the lentivirus they were initiated with. Transient over-expression of wild-type *Rbms3* in the lentiviral PCDH expression backbone was used here as a positive control. Statistical analysis was conducted using a paired T test; * = p < 0.05; **** = p < 0.0001.

In order to elucidate the mechanism(s) by which RBMS3 silencing promotes the progression of BRAF^V600E^-driven lung tumors, we considered putative mechanisms of action of RBMS3. For example, it has been reported that RBMS3 binds to the 3’ UTR of *c-MYC* mRNA such that its reduction or loss might lead to elevated expression c-MYC mRNA and protein, and thus it may serve as a possible inhibitor of WNT>β-Catenin signaling (Penkov et al. 2000; Li et al. 2011; Chen et al. 2012; Yang et al. 2017; Wu et al. 2019; Zhu et al. 2019). Previous work from a number of groups have demonstrated a critical role for WNT>β-Catenin>c-MYC signaling in normal mouse lung development and homeostasis, and also in the development of KRAS^G12D^- or BRAF^V600E^-driven lung cancers in GEM models (Desai et al. 2014; Juan et al. 2014; Kadzik et al. 2014; Tammela et al. 2017; Nabhan et al. 2018). To determine if reduced expression of RBMS3 might influence WNT>β-Catenin>c-MYC signaling in the mouse lung, we peformed immunohistochemical analysis of c-MYC expression in tumor-bearing sections of mice from *Braf*^*CAT*^; *H11*^*LSL-CAS9*^ mice initiated with either lenti-sgNT-CRE or lenti-sgRBMS3-CRE viruses (Supp. Fig. 3). This analysis revealed only a subtle increase in c-MYC expression in lung tumors that lost RBMS3. However, when similar tissue sections were analyzed for expression of β-Catenin by immunofluorescence we observed a robust increase in β-Catenin protein expression in BRAF^V600E^-driven lung tumors without RBMS3 (Fig. 5). Collectively, this analysis suggests that loss of *Rbms3* may cooperate with BRAF^V600E^ to promote lung cancer progression by increasing β-Catenin expression, thus building on what we have previously described (Juan et al. 2014) and elucidating RBMS3 as a novel component in this regulatory pathway.

**Figure 5.**
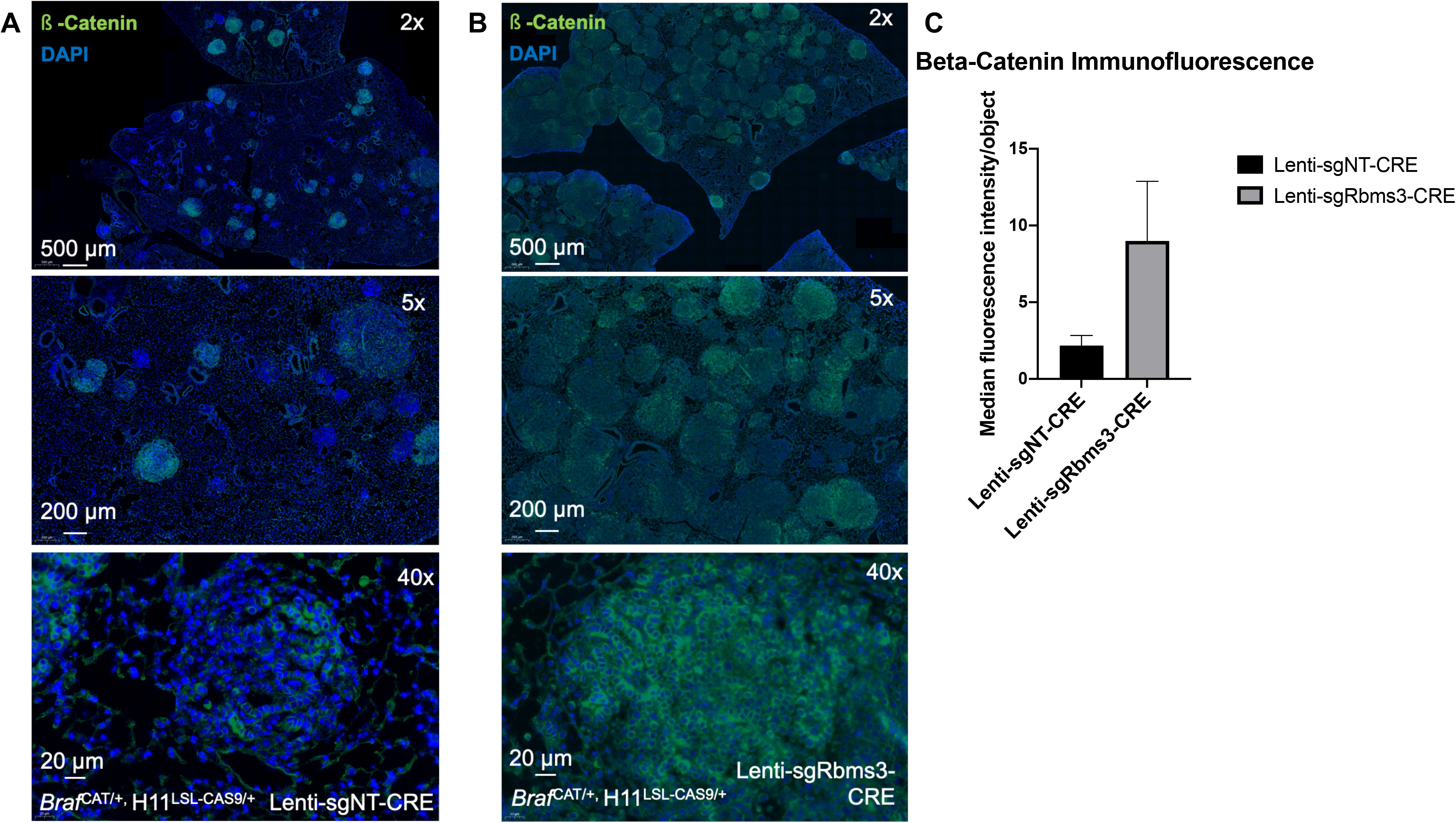
ß-Catenin protein levels are elevated in mouse lung tumors with edited *Rbms3*. (**A**) Panels of ß-Catenin immunofluorescent staining analyses in the indicated genotypes of tumor bearing FFPE mouse lung sections shown at 2x, 5x, and 40 magnifications. Scale bars are shown in white in the bottom left corner of each image. (**B**) Median fluorescence intensity quantitation using cellprofiler software. Quantitation revealed a trend of increased median staining intensity in lungs without *Rbms3*, but no statistically significant difference was seen.

To asses the possible prevalence of *RBMS3* loss in human lung cancer, we evaluated existing patient data from The Cancer Genome Atlas (TCGA) available through the cBioPortal for cancer genomics (Cerami et al. 2012; Gao et al. 2013). Initial analysis indicated that point mutations in the *RBMS3* gene are rare in this collection of human lung tumors. Analysis of changes in the region of chromosome 3p24 where *RBMS3* is located in lung cancer patients gave a striking pattern (Supp. Fig 4A). We noted that >45% of lung adenocarcinomas and >89% of lung squamous carcinomas displayed copy number alterations of *RBMS3* on chromosome 3p24 (Supp. Fig. 4B–D). However, it must be noted that the majority of these cases showed loss of the entire chromosome 3p arm, not specifically *Rbms3*. More precisely, the *RBMS3* gene; 77.8% of squamous cell carcinoma patients (87% of the patients who lost *Rbms3*) and 38.7% of adenocarcinoma patients (86% of patients who lost *Rbms3*), showed loss of the entire 3p arm (Supp Fig. 4). Furthermore, the loss of chromosome 3p24 in these patients also resulted in a worse prognosis for such lung cancer patients compared to those whose lung tumors retained chromosome 3p24 (Supp Fig. 4E). Hence, our lung cancer patient analysis revealed that the region of chromosome 3 that encompasses *RBMS3* is frequently lost in human lung cancers, and that is correlated with poorer patient prognosis.

## DISCUSSION

Genetic transposition by mobile elements, first identified by Barbara McClintock, has contributed to the identification of numerous genes that contribute to malignant transformation such as *c-MYC* and various *WNT*s in hematolgic and mammary neoplasias respectively (Hayward et al. 1981; Neel et al. 1981; Nusse and Varmus 1982). More recently, the TC1/Mariner-based *Sleeping Beauty* transposase was resurrected in conjunction with various designed *T2/Onc* transposable elements to identify tumor suppressors and oncogenes involved in cancer initiation, progression and maintenance (Dupuy et al. 2005; Copeland and Jenkins 2010; Mann et al. 2012; Mann et al. 2015). However, despite the power of such forward genetic screens to identify candidate genes involved in oncogenesis, it is important that any hits identified in such screens be separately validated. Historically, the use of retroviral systems for insertional mutagenesis has been successful only in mammary and hematopoetic tumors, limiting the scope of their potential within the majority of solid tumors. Functional screens that leverage SB transposon-mediated insertional mutagenesis have helped identify an impressive number of cancer progression-related genes in many different tumor types, including, but not limited to (Collier et al. 2005; Dupuy et al. 2005; Collier et al. 2009; Copeland and Jenkins 2010; Brett et al. 2011; Mann et al. 2015; Takeda et al. 2016; Beckmann et al. 2019; Rahrmann et al. 2019; To et al. 2021). Indeed, SB transposon mutagenesis is a powerful tool *in vivo* not just for mouse models cancer, but also drives solid tumor formation in zebrafish (McGrail et al. 2011).

In this study, we successfully used SB to identify a number of tumor suppressor genes and oncogenes that may be able to cooperate with BRAF^V600E^ in the mouse lung to drive cancer progression from benign adenomas, including: *Rbms3*, *Foxp1*, *Arid1b*, *Snd1*, *Gnaq*, and *Cux1*. Many of these genes have been previously shown to play a role in cancer. For example, inactivation of *Cux1* has previously identified as a promoter of tumor progression previously identified in a SB screen performed on myeloid malignancies (Wong et al. 2014). Of these genes, each has been previously indicated to play a role in cancer. Inactivation of *Cux1* was previously identified as a promoter of tumor progression on a SB screen performed on myeloid malignancies (Wong et al. 2014). *GNAQ* was recently shown to be an actionable target in uveal melanoma, where combining chloroquine with MEK1/2 inhibition was demomstrated to be an efficacious strategy (Truong et al. 2020). In addition to roles in cancer, *Foxp1* has specifically been shown to transcriptionally regulate lung endoderm development, but it also plays less-straight forward, context-dependent roles in different cancer types *(Koon et al. 2007; Li et al. 2016)*. Interestingly, *SND1* has been found fused to *BRAF* to form an oncogenic fusion gene in never smoker lung adenocarcinoma (Jang et al. 2015). *Rbms3* was previously identified in another SB screen in the lung, but no analysis or validation was successfully shown (Aiderus et al. 2021). Collectively, our SB screen identified a wide variety of potential lung cancer progression promoting genes.

Studies using genetically engineered mouse models that harbor either BRAF^V600E^ or KRAS^G12D^ oncogenic driver mutations have dramatically improved our understanding of tumor initiation and progression, and revealed there is still much left to learn due to the complex nature of these processes. Cancer has many steps, that involves both the activation of oncogenes, as well as the inactivation of select tumor suppressor genes over time. Expression of BRAF^V600E^ alone in the lung epithelium is sufficient to initiate multi-focal, benign adenomas of the lung that undergo a senescence-like growth arrest by 14 weeks, and these adenomas mostly fail to progress to fully malignant lung cancer without additional genetic hits (Dankort et al. 2007). Previous studies have identified a variety of genes that can cooperate with BRAF^V600E^ to form fully malignant lung cancer in the mouse, including *Tp53*, *Ink4a/arf*, *PiK3ca*, *Cdkn2a*, *c-Myc*, and *Ctnnb1 (Dankort et al. 2007; Trejo et al. 2013; Juan et al. 2014; Shai et al. 2015; van Veen et al. 2019).* Additionally, both overlapping and unique cooperating genes have been identified through characertization of genetically engineered mouse models of KRAS^G12D^ or KRAS^G12V^—driven lung cancer, including *Tp53*, *NKX2-1*, *Pten*, *APC*, and *Keap1* to name a few (Jackson et al. 2001; Sanchez-Rivera et al. 2014; Romero et al. 2017; Tammela et al. 2017). Previous studies have shown many of these genes can cooperate both with oncogenic KRAS models BRAF^V600E^ in a number of different tissues of origin (such as in pancreatic, thyroid, and melanoma models) ultimately demonstrating the relevance and importance of these cooperating genes in different tumor types (Dankort et al. 2009; Charles et al. 2011; Damsky et al. 2011; Collisson et al. 2012; Guerra and Barbacid 2013; Charles et al. 2014; McFadden et al. 2014; Mann et al. 2015). Future studies will likely reveal the relevance of *Rbms3* as a cooperating gene in the context of additional oncogenic or tissue of origin models.

*Rbms3* is a relatively newly discovered gene with lots of room to improve our understanding of its roles in normal physiology as well as its roles during tumor intitiation, maintenance, and progression. It contains two canonincal RNA binding motifs and is closely related in structure to members of the c-MYC gene single-strand binding protein family (MSSPs) (Penkov et al. 2000). Here, we have shown that *Rbms3* loss, together with the BRAF^V600E^ oncogene, can promote cancer progressionmay occur through the ability of RBMS3 to bind to the 3’ UTR of c-MYC, as well as upregulation of key components of the WNT signling pathway, specifically resulting in accumulation of higher protein levels of ß-Catenin (Penkov et al. 2000; Li et al. 2011; Liang et al. 2015; Wu et al. 2017). A number of other studies have suggested similar important roles for *Rbms3* as a tumor suppressor gene, and a regulator of tumor progression in breast cancer, esophageal squamous cell carcinoma, ovarian cancer, gastric cancer, and lung squamous cell carcinoma (Li et al. 2011; Chen et al. 2012; Liang et al. 2015; Wu et al. 2017; Yang et al. 2017; Wu et al. 2019; Zhu et al. 2019). Importantly, as shown in Supplementary Figure 6, *Rbms3* is located on a chromosomal region that clearly undergroes copy number loss in a significant number of lung cancer patients. Collectively, our data provides a strong rationale for improving our understanding of its molecular and cellular functions in conjunction with common lung cancer oncoproteins such as BRAF^V600E^.

Numerous studies have demonstrated that WNT/ß-Catenin signaling is critical in normal lung physiology, as well as in the initiation, maintenance, and progression of lung cancer (Desai et al. 2014; Juan et al. 2014; Kadzik et al. 2014; Tammela et al. 2017; Nabhan et al. 2018). The unique approaches utilized in those studies, both targeted, and objective screens, have intersected on a repeat implication to the importance of the same pathway: WNT signaling. This evidence not only emphasizes the importance of this pathway in lung cancer at different stages, buts brings about a critical question, begging whether all roads eventually lead to WNT signaling as a major player in the pathway of lung cancer progression? Our group has previously shown in the context of BRAF^V600E^-driven lung cancer, that signaling thorugh ß-Catenin is both necessary and sufficient for NSCLC, and the c-MYC is a major downstream effector of both ß-Catenin and the MAPK pathway (Juan et al. 2014). In this study, the observed role of *Rbms3* in promoting progression had only a modest effect on c-MYC protein levels, but clearly resulted in elevated levels of ß-Catenin in lung tumors. Previous work has demonstrated that growth arrest, initiated with BRAF^V600E^ signaling through the MAPK (RAF>MEK>ERK) pathway alone, can be by-passed through the constitutive activation of either c-MYC or ß-Catenin (as well as other genes such as silencing of *Tp53* or *Ink4A/ARF* or activation of PIK3CA^H1047R^ in conjunction with BRAF^V600E^ (Dankort et al. 2007; Trejo et al. 2013; Juan et al. 2014). It is possible the loss of *Rbms3* could be affecting additional pathways and acting through different mechanisms, for example, at least one other study has implicated RBMS3 in regulation Twist1 expression in metastatic breast cancer (Zhu et al. 2019). This will likely be revealed upon further analyses in future studies.

In summary, our work has revealed *Rbms3* is a novel BRAF^V600E^ cooperating gene in mouse models of lung adenocarcinoma by leveraging *SB* transposon mediated screening. Importantly, *Rbms3* is located on a region that clearly undergroes copy number loss in a significant number of lung cancer patients. This study identifies novel genetic alterations following transposon insertional mutagenesis and how they can cooperate to accelerate tumor progression in lung cancers driven by the BRAF^V600E^ oncoprotein.

## ACKNOWLEDGEMENTS

This project began in 2008 at the University of California San Francisco, but was completed at the University of Utah’s Huntsman Cancer Institute in 2019. With that said, the authors would like to thank everyone who contributed to this project over the years. Strong efforts have been made to thank, and properly acknowledge everyone who has contributed to this project over the last 11 years, but the authors sincerely apologize if anyone or anything was missed. The authors thank Dr. Shin-Heng Chiou from Dr. Monte Winslow’s lab at Stanford University for assistance in preparing reagents and executing experiments for these studies. The authors thank Dr. Trudy Oliver’s lab, especially Dr. Rachelle Olsen and Dr. Gurkan Mollaoglu at the Huntsman Cancer Institute for their assistance in providing proper controls and troubleshooting of experiments performed for this manuscript. The authors thank Dr. Eric Snyder’s lab at the Huntsman Cancer Institute, especially Dr. Soledad Camolotto, for technical expertise and assistance supporting these studies. The authors thank the Huntsman Cancer Institute in Salt Lake City, UT for the use of the following shared resources: (1) The University of Utah Health Science Center DNA Sequencing Core Facility. (2) The University of Utah Health Science Center Flow Cytometry Core Facility (and the National Cancer Institute through Award Number 5P30CA042014-24). (3) The Biorepository and Molecular Pathology Shared Resource, specifically research histology (operated by ARUP labs), and nucleic acid isolation.

## AUTHOR CONTRIBUTIONS

This project was initiated by J.J. and M.M. in 2011 and was brought to fruition by A.V. and M.M. at this time. A.V., J.J. and M.M. designed and executed experiments, interpreted data in collaboration with S.C., K.M.M., J.Y.N., A.R., D.A., A.G., and M.B.M. wrote the manuscript and shepherded it through review. M.T.S. and J.E.V. assisted with *in vitro* and *in vivo* experiments. C.S.H. and W.A.W. assisted with generation of critical reagents. A.G. assisted with pathological evaluation of tumors. L.v.d.W performed the linkage-mediated PCR. M.B.M., J.Y.N., A.R., and D.J.A. assisted with sequencing and computational analyses. C.S. helped with data mining of cBioPortal. A.L. performed mouse genotyping-related tasks.

## MATERIALS AND METHODS

### SB tumor sequencing, informatics and statistical analyses

#### Illumina sequencing

Tumor DNA was extracted from formalin-fixed paraffin-embedded (FFPE) tissues using the Gentra PureGene cell kit (Qiagen; 158767) according to the manufacturer’s instructions, and barcoded genomic fragments containing transposon-genome junctiojns sequences were amplified using linker-mediated PCR (LM-PCR). Illumina sequening was previously described [Uren et al., 2009]. These products were sequenced on the 454 platform, from which unique sequencing reads were generated.

#### Raw processing of sequence data

28 tumors were taken from 10 mice and pair-end sequencing was performed using customized baits and aligned to the mouse genome reference assembly GRcm38 using BWA (version 1.16). The GATK ‘indel realigner’ was used to realign reads near indels from the Mouse Genome Programme to improve indel/SNP identification. The BAM files were re-sorted to recalibrate quality scores with the GATK ‘TableRecalibration” tool. SAMtools ‘calmd” was utilized to recalculate the MD/NM tags within the BAM files. Every lane from the same library were merged into a single BAM files using Picard tools (version 1.72) and PCR duplicates were marked using Picard ‘MarkDuplicates’.

#### Merging and Filtering

The Bam files were processed using RetroSeq 9 (version 1.41) to identify pair reads where one read aligned to the reference mouse genome and the other read to the *Sleeping Beauty* (SB) transposon sequence. (Retroseq was operated in “Discovery” mode using the default parameters: Min anchor quality; 20; Min percent identity: 80; Min length for hit: 36). This generated a total of 72,981 individual putative transposon insertion regions (70 shared across all 28 tumor samples.) This sequence and analysis method does not allow the exact SB insertion sites to be identified to the resolution of genomic basepairs hence the location of transposons are referred to as insetion regions (IRs).

Overlapping, individual insertion regions (IRs) within each sample were merged using bedtools to generate a set of 41,152 IRs. Chromosome four is the donor chromosome for the TG.6070 transposon line. To reduce the effects of local-hopping that can skew the downstream statistical analysis, all IRs that were located on chromosome four (4,609) were thereby excluded from the analysis. Insertions on the other secondary scaffolds e.x. GL45693, were also excluded. This left a total of 36,510 IRs.

Further filtering of the IRs were performed by removing IRs within the regions of two known genes into which the SB concatemer preferentially inserts (on GRCm38:En2; chr5;2816569628173612 and *Foxf2*; chr13;31625816-31631403.) Nineteen genomic regions reported into which the SB transposon inserts under no selection pressure were also used to exclude IRs that are likely not to be cancer drivers. Following these final filtering step resulted in 36,426 IRs.

#### Common insertion sites and SB trunk driver analysis

To identify common insertion regions (CIRs), where the SB transposon inserts with a greater chance than compared to random, a Gaussian kernel convolution (GKC) statistical framework was used as described previously (Takeda et al. 2016; Newberg et al. 2018b). Briefly, to detect CIRs using an IR bed file, a GKC method (March et al. 2011) was employed using 15,000, 30,000, 50,000, 75,000, 120,000, and 240,000 kernel widths. When CIRs were detected over several kernel widths, the CIRs were merged and the smallest kernel width is reported as representiative CIR. For highly significant CIRs with narrow spatial distributions of insertion regions, the 15K kernel is typically the scale on which CIRs are identified.

A separate bioinformatics workflow, SBseq (Mann et al. 2016) was used to map SB insertions to the mouse genome used a custom bioinformatics workflow modeled to perform similarly to the analysis pipeline reported previously (Mann et al. 2015). The SBseq workflow used the subset of IR reads that permited the detection of unique, directional SB insertions with base pair resolution at TA-dinucleotides. First, the BAM files were processed to map the short reads to identify a sub-set of the original data that was able to detect single nucleotide mapping resolution and the orientation of the SB insertions with respect to the direction of gene expression. Briefly, we converted from BAM format to FASTQ format using SamTools and then mapped the FASTQ reads to unique mouse TA dinucleotide sites. Next, TA sites were converted to BED format, and masking of known SB insertion hotspots (as above for CIR), was performed. A final BED formatted file contained 6,396 non-redundant SB insertions at single nucleotide resolution, including donor chromosome 4 insertions, from 28 lung masses. SB Driver Analysis (http://sbcddb.moffitt.org/software/) was performed as previously described (Newberg et al. 2018a) using bed formatted files generated by the RetroSeq and SBseq bioinformatics pipelines, to generate Progression and Trunk Driver (with read depth cut-off: 3) candiate genes, for all chromosome excluding the SB transgene donor chromosome 4.

A gene-centric statistical method was used to identify CIS genes, and genes that had 5 or more read counts, and had insertions in three or more tumors were selected as trunk driver genes. A Bonferroni correction was added to help eliminate false positives, and adjust the p-values (Brett et al. 2011). This initial list of candidate genes was further analyzed by bioinformatics tools such as Ingenuity Pathway Analysis, as well as STRING and DAVID tools to assess biological relevance, followed by cross-referencing of human tumors analyzed by TCGA.

### Animals

All mice were housed in an environmentally controlled room, and all animal care and experimental procedures were approved by (and in accordance with) the Institutional Animal Care and Use Committee (IACUC) Office of the University of California San Francisco, and also later on during this project at the Huntsman Cancer Insitute at the University of Utah. Genetically engineered mouse breeding and genotyping was conducted as previously described (van Veen et al. 2019). The BRAF^CAT^ mouse was previously described, and the H11^LSL-CAS9^ mouse was kindly provided from Monte Winslow’s lab at Stanford University (Chiou et al. 2015; van Veen et al. 2019).

All viruses were administed in a Biosafety Level 2+ room, as is regulated by the Institutional Biosafety Committee Guidelines. Adeno-CRE virus (University of Iowa) was delivered through nasal instillation, and Lenti-CRE virus (described in detail below) was delivered through direct intratracheal instillation under isoflurane anesthesia. The *T2/Onc2* (Strain 6070 [B6;C3H TgTn(sb-T2/Onc2)6070Njen] (MGI: 3613048) was bred and genotyped as previously described (Dupuy et al. 2005)Adeno-CMV-CRE virus was purchased from Viraquest (North Liberty, Iowa). In conjunction with IACUC policy, mice were euthanized when the animal demonstrated any serious health concerns or signs of suffering. The Ullmann-Cullere Body Conditioning Score (BCS) was used to determine if euthanasia endpoints were necessary (Ullman-Cullere and Foltz 1999).

Upon euthanasia, mouse lungs were inflated using either PBS or 10% neutral buffered formalin for perfusion through the larynx, followed by an additional cardiac perfusion of the lung though the right ventricle of the heart until the lungs turned white. Lungs were fixed for 24 hours in formalin prior to transfer to ethanol for paraffin-embedding and sectioning at 4 microns.

### Generation of Rosa-LSL-CAGGS-SB11 mice

The *Rosa*26-CAGGSCAGGS-*lox*P-STOP-*lox*P (64)::*SB* mouse was created by taking a *Rosa26* targeting vector (Collier et al. 2005; Dupuy et al. 2005) and engineered as follows: A construct with an EcoRV restriction site followed by 521bp of homology to the *Rosa26* locus, intron 1 and exon 2 of the mouse Engrailed 2 gene, the CAGGS promoter, and loxP-flanked EGFP, 2xSV40 polyA sequences, and a BGH polyA sequence was modified by inserting a linker containing NotI, XhoI, and SphI restriction sites into a SalI restriction site downstream of the 3’ loxP site. pCMV-SB11 (Addgene plasmid # 26552, a gift from Dr. Perry Hackett, University of Minnesota) was modified to include a NotI site downstream of the SB11 CDS. This vector was digested with EagI and NotI and ligated into the NotI site of the modified *Rosa26* targeting vector. A sequence containing SV40 polyA, a flippase-recognition target (FRT), PGK promoter, neomycin resistance cassette and BGH polyA signal, a second FRT, and 601bp of homology to the *Rosa26* locus was isolated from the initial Rosa targeting vector and ligated into the XhoI site downstream of SB11. This shorter targeting vector was then recombined into a larger *Rosa26* targeting construct containing 3.5kb and 2.9kb of Rosa26 homology on the 5’ and 3’ ends, respectively. This plasmid was linearized and transfected into E14 129/Ola mouse embryonic stem (ES) cells. DNA was isolated from selected ES cell clones, digested with ApaI, and screened by Southern blot using a probe outside of the targeting construct to identify clones with restriction fragment length polymorphismswith indicating integration of the CAGGS-LSL-SB11 cassette. One clone (A3) was injected into blastocysts to generate 13 chimeric mice. Chimeric males were mated to 129/SvJ females; one chimera was able to propagate the targeted allele through the germline.

Southern blot probe sequence:

>probe us1

Ctgggaaggttccttaagaagttatgttctgagaccattctcagtggctcaacaacacttggtcaaaaattttaattctcccctcagagaaatggagtagttactccactttcaagttccttataagcttaccatcaaccttatagtacactctagatgtcctgaaatatttctatcagaacaaggtagtataaagctggtaggtatacaaaacgctagactagtttctatccctgacccttaatctgctagtatatccgtaggaagttgcttaagtgccactagtacca.

### Cell lines, 2D and 3D culture conditions, and imaging

HEK-293T cells were maintained in DMEM media supplemented with 10% fetal bovine serum and 1% penicillin/ plus streptomycin. HEK-293T cells used for these studies have been authenticated by STR profiling and mycoplasma testing is done quarterly using PlasmoTest (Invivogen).

Organoids were established by dissociating lung tissues minced with scissors in digestive media comprised of collagenase (400 U/ml), dispase (5 U/ml), elastase (4 U/ml), and DNAseI (0.25 mg/ml) in Advanced DMEM:F12 HAM media in a 37°C shaker for 30 minutes. After generating a single cell suspension, cells were strained using 100, 70, and 40 micron filters. Red blood cell (RBC) lysis was performed at room temperature by incubating each sample with 1x RBC Lysis Buffer (eBioscience; 00-4333-57). Finally, cells were seeded at 50,000 cells/well in matrigel (Corning; #356327) in a 24-well plate. Organoids were grown in organoid culture media containing Advanced DMEM/F12, 1x B-27 (Thermofisher; #17504001), 1x N-2 (Thermofisher; #17502001), 1% Penicillin/Streptomycin, 1.25 mM N-acetylcysteine (Sigma-Aldrich; #A0737), 10 nM Gastrin (Sigma-Aldrich; G9020), 10 μM Nicotinamide (Sigma-Aldrich; #47865-U), 100 ng/mL R-Spondin-1 (Peprotech; #315-32), and 100 ng/mL Noggin (Peprotech; #25038) (Boj et al. 2015; Zewdu et al. 2021). To enrich for cells positive for constitutive BRAF^V600E^ signaling, no ligands that could activate MAPK signaling (i.e. EGF or FGF) were added in this organoid media recipe. All phase contrast images of cultured cells were taken with an EVOS microscope at 10x magnification (Invitrogen), and contain a scale bar of 400 microns in the image.

### Immunohistochemistry and immunofluorescence of lung sections

Immunohistochemistry was performed as previously described, with the rabbit primary antibody against c-MYC (Santa Cruz; sc-764; 1:150) (Mollaoglu et al. 2017). Similarly, immunofluorescence staining of the Sleeping Beauty Transposase was performed by fixing mouse lungs in zinc-buffered formalin, processed and embedded in parrafin, cut into 5 micron sections, and mounted on glass slides. Citrate mediated antigen retrieval was peformed, followed by staining with the indicated primary antibody from R&D Systems.

### Slide scanning, imaging, and histological analyses and quantification

Hematoxylin and eosin (H&E) stained slides, and immunofluorescence slides from the SB screen shown in Figure 1 were scanned using an Aperio Scanscope Scanner. Following H&E staining of sectioned lungs from remaining figures, as well as c-MYC immunohistochemistry, or immunofluorescence analysis with ß-Catenin, slides of sectioned mouse lungs from each indicated genotype were loaded and scanned automatically by the 3D Histech Pannoramic MIDI scanner (Thermo Fisher). Slides were imaged and analyzed using Caseviewer Software or QuantCenter analytical center provided on the 3D Histech Slide Scanner. Tumor burden was manually calculated on each lung lobe and total tumor area was compared to total lung area. Tumor diameters were measured using QuantCenter software from 3D Histech. Cellprofiler was used to quantitate median fluorescence intensity with a previously described pipeline following immunofluorescence analysis of mouse tumor bearing lungs (van Veen et al. 2019).

### Plasmid cloning, lentivirus production, cell transduction

The CAG-HA-RBMS3-PCDH cDNA expression plasmid was cloned using a cDNA template made from RNA from the lungs of a wild-type mouse using the Q5 polymerase (NEB) and restriction endonuclease cloning with the following primers: 5’: tttttGAATTCCCACCATGTACCCCTATGATGTGCCAGACTACGCCGGCAAACGCCTGGATCAGCCA CAA. 3’: TttttgcggccgcCTATGGTTTGGACTGTTGGAAGGA. EcoRI and NotI enzyme were used to insert the cDNA of *Rbms3* into the mammalian expression plasmid PCDH. Lenti-sgNt/CRE and lenti-sgLkb1/CRE was a gift from Monte Winslow (Addgene plasmid # 66894) (Chiou et al. 2015). Three sgRNAs designed against *Rbms3* were cloned into the pLL3.3 sgRNA-CRE vector by modifying the original sgLKB1 plasmid with the Q5 Site-Directed Mutagenesis kit (NEB). The following sgRNAs against *Rbms3* were used and pooled to make 2 lentiviruses.

Pool 1: 1).GTACACGTACTACTGTCCTC. 2). GAGCACGTCATGGACGCCAC. 3). ATGCAGCCAACTAACATCGT.

Pool 2: 3). ATGCAGCCAACTAACATCGT. 4) TTGGACACGTGATATCCACC. 5). ATCAAGCTATGTCAACCGTA.

Successful clones were verified by Sanger sequencing. Lentiviral supernatants were generated by co-transfection of HEK-293T cells using Transit-X2 (Mirus; #MIR 6004) with a 3-vector lentiviral system: using either the non-targeting sgRNA expression vector or sgRBMS3 sgRNA pool 1 or pool 2 combined with the lentiviral packaging and envelope plasmids PCMV-Δ8.9 and PCMV-VSVG. pCMV-VSV-G was a gift from Bob Weinberg (Addgene plasmid # 8454). Virus was collected 36, 48, 60, and 72 hours post-transfection, and filtered using a 45 micron filter. Viral supernatants were concentrated by centrifugation at 25,000 rpm for 105 minutes at 4°C. Viral pellets were resuspended in 1x PBS, and stored at −80C for long-term, storage. Viral titering was performed using KP1 cells, and flow cytometric analysis of RFP+ cells as previously described (Camolotto et al. 2018).

### DNA isolation and the Surveyor assay

Lung tumour tissue was microdissected and isolated from FFPE tumor blocks and DNA was purified using the QIAamp kit (Qiagen; catalog #56404). Alternatively, DNA was isolated from cell lines using the DNeasy Blood and Tissue kit (Qiagen; catalog #69504). The surveyor assay mutation detection kit was used according to the manufacter’s instructions (IDT; product number 706025). PCR amplicons of *Rbms3* for the surveyor assay were created with the following primers: 5’ CTGGATCAGCCACAAATGTACCCCC. 3’ TGCTCTGGACCTGGTATGT. The following PCR conditions were utilized with the Q5 polymerase according to the manufacturer’s instructions for 25 or 50 μl reactions (NEB): 98° for 30s, 32 cycles of (98° for 10s, 53° for 20s, 72° for 40s), 72° for 2 m, and then stored at 4°C for short periods or −20°C for long-term storage.

### RNA isolation and qRT-PCR

Tumour tissue was microdissected and isolated from formalin-fixed paraffin embedded (FFPE) blocks and RNA purified using the RNeasy FFPE Kit (Qiagen; catalog #73504). RNA was purified from cultured organoids following dissociation with TrypLE (Thermo Fisher), pelleting, and resuspending the organoid cell suspension in Trizol. One-fifth volume of chloroform was added, and the tube was shaken vigorously, followed by centrifugation for 15 minutes at 12,000 x g at 4°C. The aqueous phase was transferred to a new tube, and 10 μg of glycogen (Thermo Fisher; #R0551) was added. RNA was precipitated with 1/10 volume pH 5.2 3M Sodium Acetate (pH 5.2; Thermofisher; catalog #R1181) and 0.5 mL of isopropanol. After mixing the tube was incubated at −80°C for 30 minutes. The mixture was then centrifuged for 10 minutes at 12,000 x g at 4°C, the supernatant removed, and the pellet was washed with 1 mL of cold 75% ethanol. After vortexing, the samples were centrifuged again, before the pellets were air dried, and resuspended in RNase-free water.

cDNA template was generated using 250 ng of RNA with the iSCRIPT reverse transcription supermix (Bio-Rad; catalog # 1708841) according to the manufacter’s recommended protocol. SSOAdvanced Universal Probes Supermix (Bio-Rad; catalog # 1725280) was used also according to the manufacter’s protocol. qRT-PCR was performed using Taqman Gene Expression assays (Applied Biosystems; Thermofiosher) and the following 20x probes: *Ppia* (mm03302254_g1) as a housekeeping gene for normalization, and *Rbms3* (mm01350499_m1; mm00618362_m1; mm01350496_m1).

### cBioPortal Analysis

Point mutations were defined as single base-pair alterations and copy number alterations were defined with copy number values less than or equal to −1, consistent with TCGA standards. Search criteria involved listing chromosome arm 3p as a separate field that was queried and automatically aggregated. Individual gene CNA analysis was conducted through downloading individual patient .cnv files and aggregating manually. Point mutations for individual genes were queried directly within the site interface. Copy number variation (CNV) data were collected via cBioPortal for TCGA-LuSC (n=487) and –LuAD (n=500) projects from the PanCancer Atlas. From these data, deletions were defined as a copy number equal to −1 and gains defined as copy number equal to 1. Using these thresholds, it was shown that in TCGA-LUAD and –LUSC, respectively, the chromosome 3p arm was deleted in 77.8% and 38.7% of patients. For RBMS3, specifically, deletion occurred in 45.4% and 89.2% of patients in TCGA-LUAD and –LUSC, respectively. Kaplan-Meier curves were generated between cohorts that incurred a deletion of the 3p arm against those that did not incur a deletion.

## Supplemental Figure legends

**Supplemental Figure 1.**
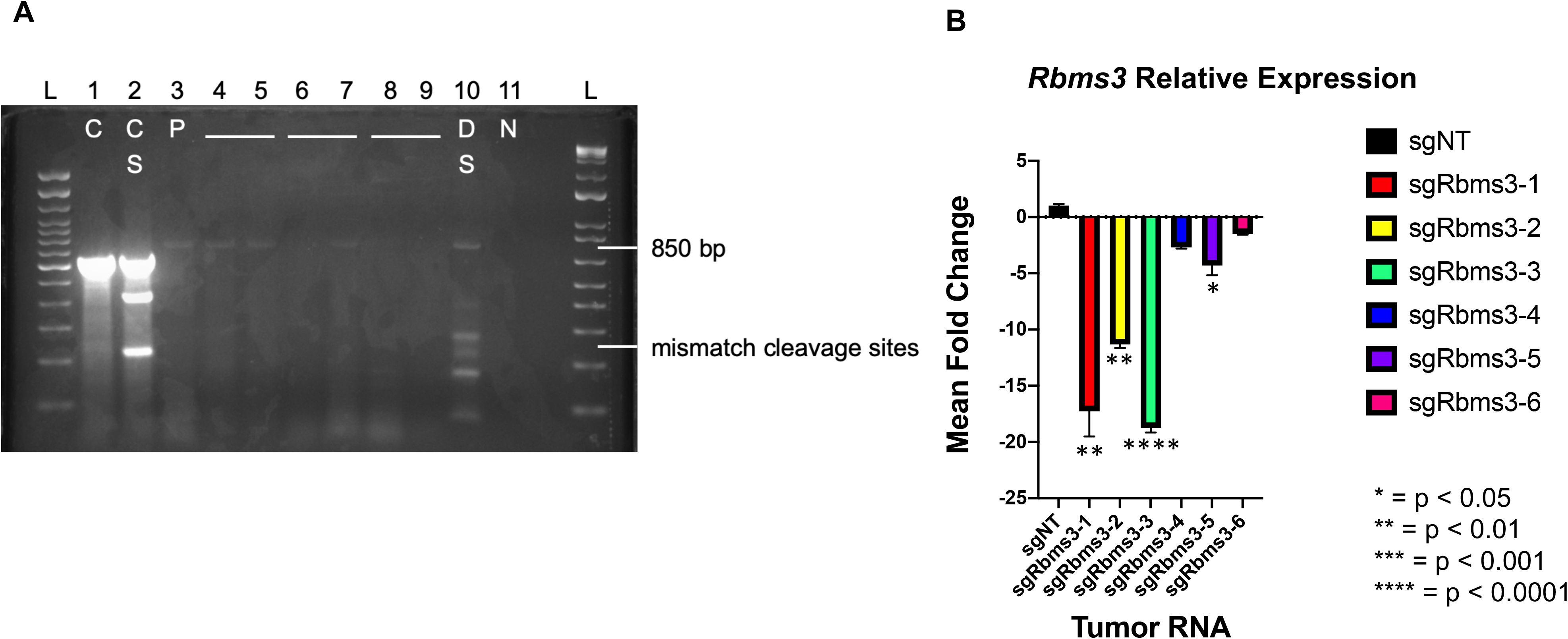
Validation of *Rbms3* loss in tumors initiated with sgRbms3-CRE lentivirus. (**A**) A PCR of *Rbms3* genomic locus is 850 bp, and contains each sgRNA used against the mouse *Rbms3* gene. Digestion of wild-type or mutant duplexed DNA with Surveyor enzyme reveals *Rbms3* locus is edited in large tumor DNA via CRISPR/CAS9-mediated editing: indicated by mismatch cleavage fragments below the 850 bp band in lane 10. L represents ladder. Numbers represent lanes. C is technical control (lane 1), S represents addition of Surveyor enzyme (lane 2), N is negative control (lane 11), P is positive or Wild-type *Rbms3* gDNA (lanes 3 and 4), sgRbms3 tumor gDNA (lanes 5-8)and D represents wild-type and sgRbms3 tumor gDNA duplexed with the Surveyor enzyme. (**B**) qRT-PCR indicates significantly lower *Rbms3* gene expression in 4 of 6 tumors excised from lung sections of mice initiated with sgRbms3 virus compared to those initiated with sgNT virus by a paired T-test. * = p < 0.05; **= p < 0.01; **** = p < 0.0001.

**Supplemental Figure 2.**
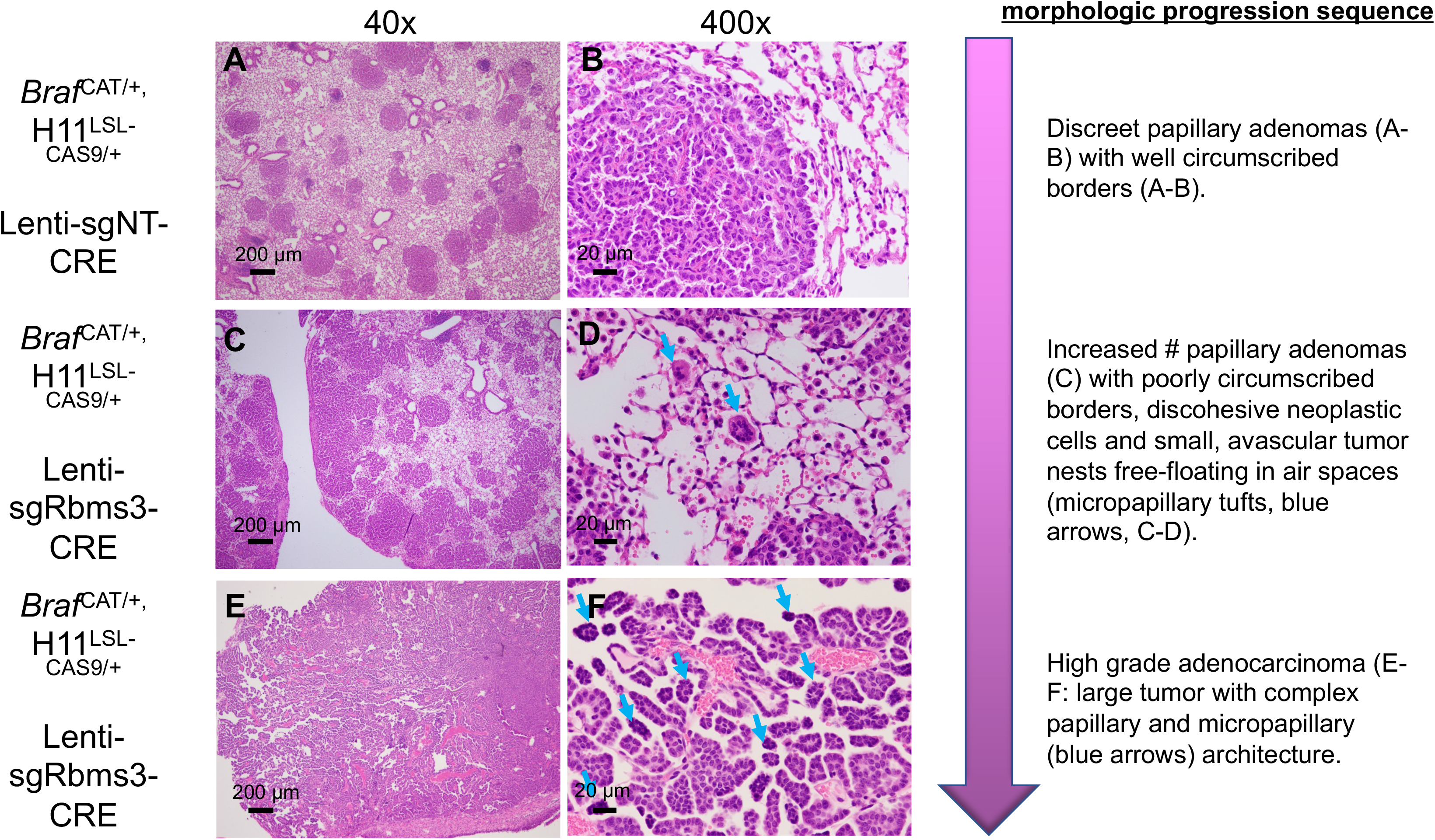
Histological analyses of the progression of grade of tumor bearing mouse lungs expressing BRAF^V600E^ with or without *Rbms3*. **(A-I)** High magnification images of formalin-fixed paraffin-embedden (FFPE) sections of mouse lungs stained with hematoxylin and eosin (H&E). **(A-B)** Tumor bearing lungs from BRAF^CAT/+,^ H11^LSL-CAS9/+^ mice initiated with sgNT-CRE virus harvested at 13 weeks post-initiation using **(A)** 40x or **(B)** 400X magnification. Shown are typical BRAF^V600E^ only papillary adenomas with well-circumscribed borders. **(C-D)** Tumor bearing lungs from BRAF^CAT/+,^ H11^LSL-CAS9/+^ mice 13 weeks post-initiation with sgRbms3-CRE virus harvested at 13 weeks using **(C)** 40x or **(D)** 400X magnification. Shown are an increased number of papillary adenomas with poorly circumscribed borders, discohesive neoplastic cells, and small, avascular nests free-floating in air spaces. Blue arrows indicate micropapillary tufts. **(E-F)** Largest tumor bearing lungs from BRAF^CAT/+,^ H11^LSL-CAS9/+^ mice 13 weeks post-initiation with sgRbms3-CRE virus harvested at 13 weeks using **(E)** 40x or **(F)** 400X. Shown here are higher grade adenocarcinoma in a larger tumor with blue arrows indicating complex papillary and micropapillary architecture. The large tumor shown here was validated to have edited *Rbms3* at the DNA and RNA level in Supplemental Figure 2. Scale bars are shown in black in the bottom left corner and are 200 microns (**A**, **C**, **E**) and 20 microns (**B**, **D**, **F**), respectively.

**Supplemental Figure 3.**
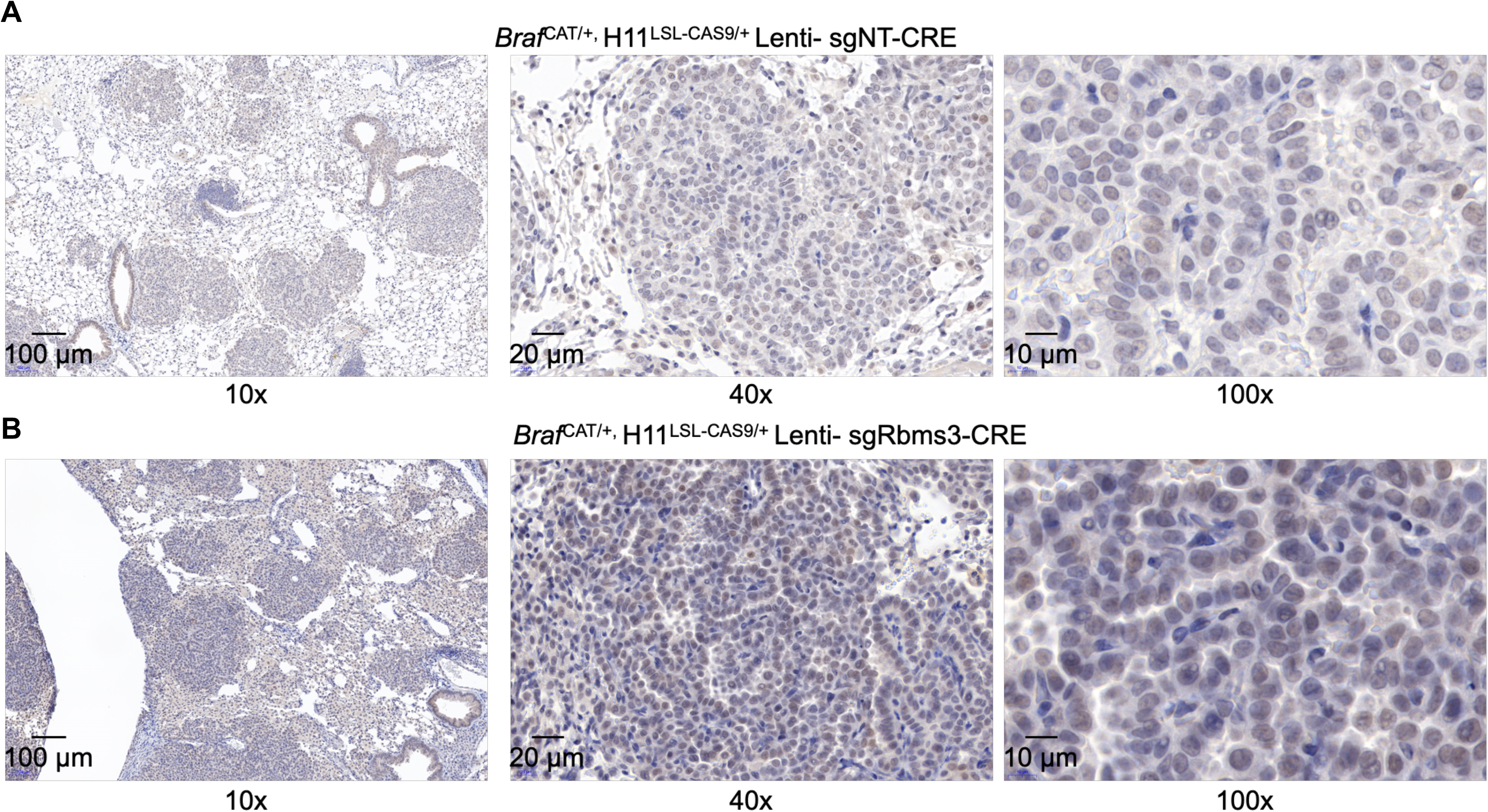
Immunohistochemistry of c-MYC protein expression in BRAF^V600E^+ mouse lung tumors with and without RBMS3. (**A**) Panels of c-MYC immunohistochemistry in BRAF^CAT/+;^ H11^LSL-CAS9^ mice initiated with sgNT-CRE at 10, 40, and 100x magnifications. (**B**) Panels of c-MYC immunohistochemistry in BRAF^CAT/+;^ H11^LSL-CAS9^ mice initiated with sgRbms3-CRE at 10, 40, and 100x magnifications. Scale bars are indicated in the bottom left corner of each image.

**Supplemental Figure 4.**
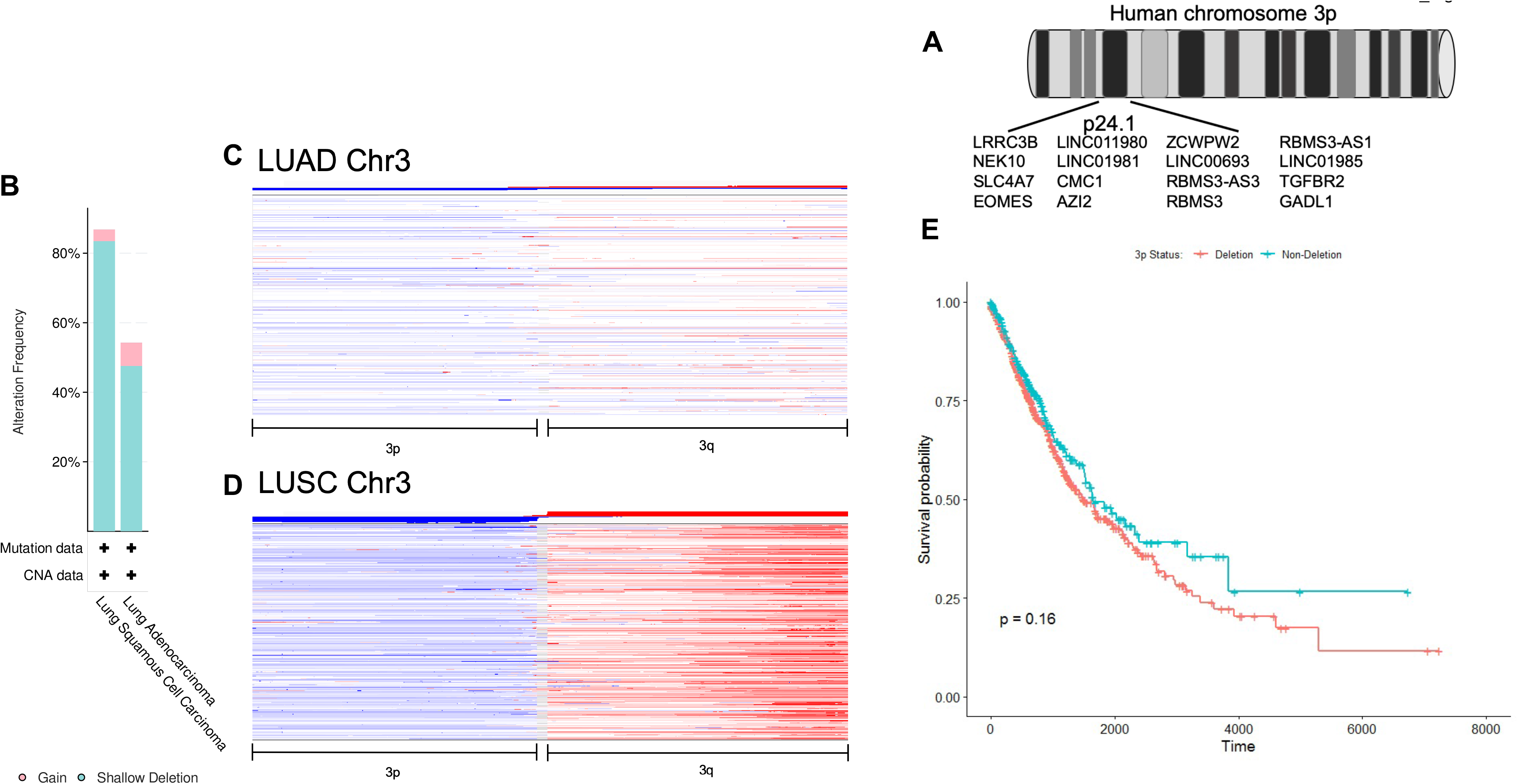
*Rbms3* is lost frequently in human non-small cell lung cancer patients. (**A**) Schematic depicting genes located on chromosome 3p24.1 (not drawn to scale). (**B**) Quantitation of cBioportal analyses of TCGA human lung adenocarcinoma (LUAD) and lung squamous cell carcinoma (LUSC) patients with identified relative copy number alteration frequences of chromosome 3p either by gain (pink) or deletion (blue). **(C) and (D)** Copy number alteration heat map depicting chromosome 3 gain/loss in red/blue (respectively) on both 3p and 3q in **(C)** lung adenocarcinoma (LUAD) or **(D)** lung squamous cell carcinoma (LUSC) patients, respectively. Each row corresponds to a patient. **(E)** Kaplain-meier survival curve of patient cohorts that do (orange) or do not (blue) have deletion of chromosome 3p. Deletion was defined to be a copy number threshold of −1 within cBioPortal data. Decreased survival is observed in patients that have lost chromosome 3p, but this was not found to be statistically significant.

